# FAVA: High-quality functional association networks inferred from scRNA-seq and proteomics data

**DOI:** 10.1101/2022.07.06.499022

**Authors:** Mikaela Koutrouli, Pau Piera Líndez, Katerina Nastou, Robbin Bouwmeester, Simon Rasmussen, Lennart Martens, Lars Juhl Jensen

## Abstract

Protein networks are commonly used for understanding how proteins interact. However, they are typically biased by data availability, favoring well-studied proteins with more interactions. To uncover functions of understudied proteins, we must use data that are not affected by this literature bias, such as single-cell RNA-seq and proteomics. Due to data sparseness and redundancy, co-expression analysis becomes complex. To address this, we have developed FAVA (Functional Associations using Variational Autoencoders), which compresses high-dimensional data into a low-dimensional space. FAVA infers networks from high-dimensional omics data with much higher accuracy than existing methods, across a diverse collection of real as well as simulated datasets. FAVA can process large datasets with over 0.5 million conditions and has predicted 4,210 interactions between 1,039 understudied proteins. Our findings showcase FAVA’s capability to offer novel perspectives on protein interactions. FAVA functions within the scverse ecosystem, employing AnnData as its input source.

## Background

Networks of physical and functional interactions among genes/proteins are widely used to understand the inner workings of cells and to visualize results from omics data, e.g., as obtained from transcriptomics and proteomics experiments. Unfortunately, most research is focused on the same 10% of human protein-coding genes (1–3). Thus, networks derived from the biomedical literature — whether through manual annotation or through automatic text mining — are heavily biased by this skewed availability of data. Networks obtained from databases such as STRING (4) consequently have many interactions for well-studied proteins. However, there are only very few interactions for understudied proteins, which are arguably the most interesting targets for network-based function prediction (3,5).

Recent publications highlight the need for new information about understudied proteins (5,6). These proteins have been largely ignored due to the Matthew effect in which proteins with proven disease importance are the subject of most studies (5). To create networks that also provide interactions for understudied proteins, one must focus on systematic high-throughput data, as these are inherently unaffected by literature bias.

One approach to address this is to use experimental approaches to systematically capture new information on, for example, physical protein interactions of understudied proteins (7). The hu.MAP 2.0 resource (8) is an example of this, providing protein complexes derived from over 15,000 proteomic experiments. By capturing physical interactions between proteins in a systematic manner, without literature bias, it provides protein interaction information for many completely uncharacterized proteins.

Complementary to this, technological advances such as RNA and mass spectrometry-based proteomics have enabled the discovery of protein functions and protein–protein relationships with greater accuracy on a large scale. Co-expression network analysis is one popular approach for this; it works by linking genes that show similar expression patterns across many samples (9,10). Many computational methods have been developed for Gene Co-expression Network (GCN) construction from bulk RNA-seq data (11), such as WGCNA (12). However, other types of data with higher resolution, such as single-cell RNA-seq (scRNA-seq) data, may allow better GCNs to be constructed. ScRNA-seq provides unbiased data on gene expression at the level of individual cells, thus capturing differences between both cell types and cell states. However, methods optimized for bulk RNA-seq may not be successful in data produced by this technology due to its sparse and redundant nature.

As high-dimensional, sparse omics data are becoming increasingly common, in part due to single-cell technologies, new methods designed to handle such data are needed. Pearson correlation coefficient (PCC) is an often-utilized technique to establish GCNs. Recently, more sophisticated computational methods have been developed for GCN construction specifically from scRNA-seq data. Some of the state-of-the-art methods are the hdWGCNA (13) and the scLink (14). hdWGCNA makes use of the WGCNA (12) protocol to construct gene networks and to identify modules with specific provisions for high-dimensionality datasets. To enable this process in scRNA-seq data, hdWGCNA identifies similar cells, which are pooled, thereby dealing with sparsity. scLink (14) manages sparsity by not considering all samples for every pair of genes, but instead focusing on the cells in which both genes are accurately measured. Additionally, the authors recommend creating GCNs with scLink using only the top 500 most highly expressed genes.

While single-cell RNA expression data provide great detail, the correlation between RNA and protein levels is far from perfect (15). It thus makes sense to consider also using proteomics data for creating GCNs, although proteomics data at single-cell resolution currently remains a rarity (16) with small numbers of cells. ScRNA-seq and proteomics can thus be viewed as two complementary types of data that can be used as starting points for predicting functional interactions, also for understudied proteins. Unfortunately, there are no computational methods designed to handle both types of data, taking advantage of their complementary nature. The biggest challenge remains that both scRNA-seq and proteomics datasets are very sparse (i.e. many transcripts/proteins are not observed in each cell/sample) and have high redundancy (i.e., many similar samples/cells are analyzed), both of which present problems for most analysis methods.

Dimensionality reduction can help address both sparsity and redundancy in data. By compressing the information into a lower dimensional space, sparsity is eliminated by combining data from across multiple cells/samples. Redundancy is also inherently reduced, because data compression is specifically achieved by not storing the same information multiple times (17). One can thus expect that applying dimensionality reduction to scRNA-seq and proteomics data provides a latent representation that is better suited for co-expression-based prediction of interactions than the original data matrices.

Dozens of dimensionality-reduction algorithms exist, which can be used to produce this latent space. These span linear methods such as truncated singular value decomposition (18,19), nonlinear transformations such as Uniform Manifold Approximation and Projection (UMAP) (20), and deep generative models such as variational autoencoders (VAEs) (21). VAEs use two deep neural networks — an encoder and a decoder (as shown in **Figure 1**) — to learn a representation from complex data without supervision (22). The encoder learns the input data and projects it into a normally distributed latent representation, parameterized by the mu (μ) and sigma (σ) layers, while the decoder attempts to reconstruct the input data from the instances sampled from the latent representation distributions. The encoder–decoder networks are simultaneously optimized to reconstruct the input data and regularize the latent representations. This ensures that the constructed latent representations follow Gaussian distributions, making them easier to interpret than those of other autoencoders, such as the tones described in (23). Unlike, e.g., singular value decomposition, VAEs can capture both linear and nonlinear relationships between the cells/samples in scRNA-seq and proteomics data. As a result, VAEs have become popular in the field of expression data (e.g., for normalization and visualization of scRNA-seq).

**Figure 1.**
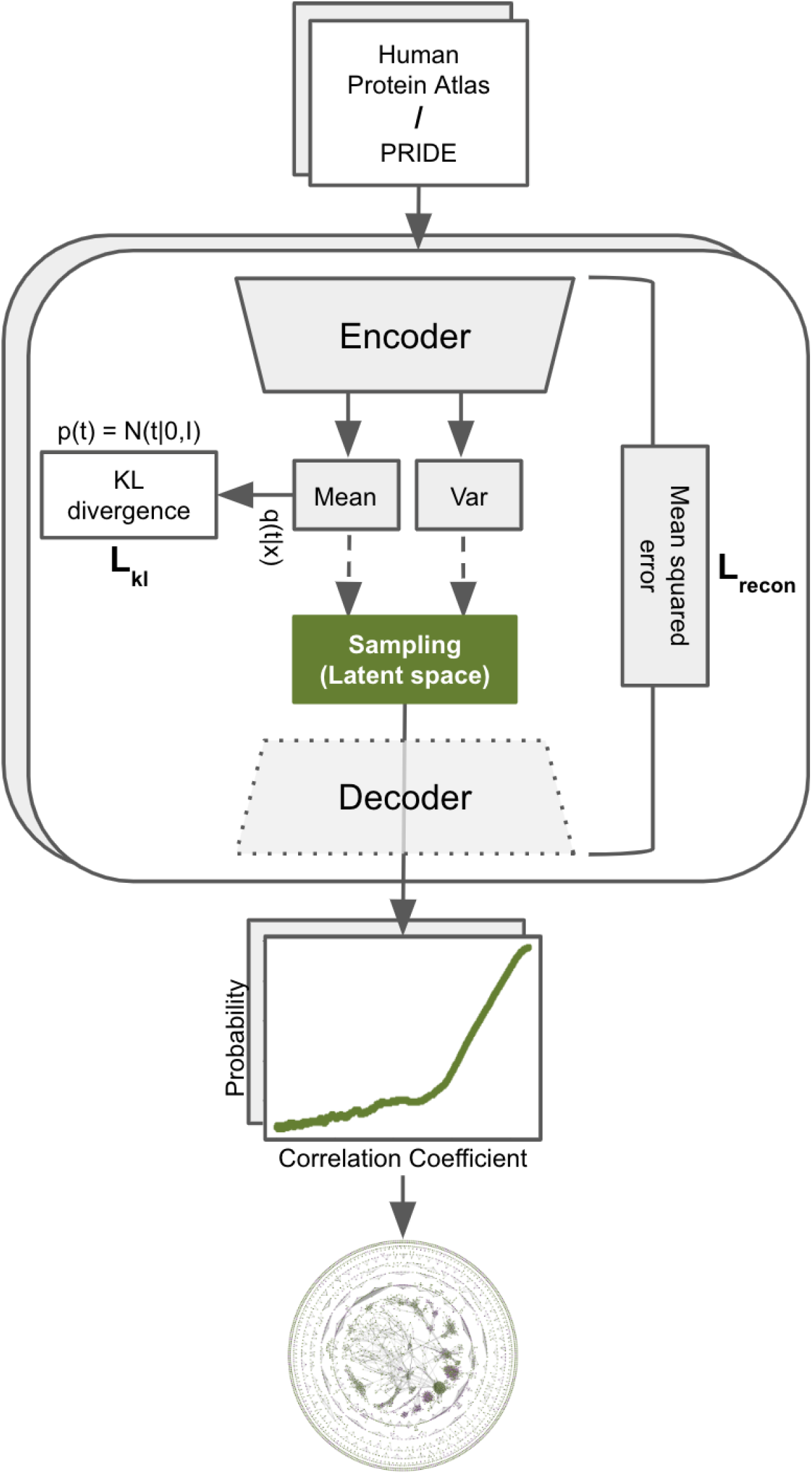
Overview of the FAVA method: from expression matrices to a combined network. The first step is to pre-process each input matrix before it is fed to a variational autoencoder (VAE). After pre-processing, the input matrix is used to train the VAE model, from which we obtain the latent space from the bottleneck layer. In this latent space, we calculate the Pearson correlation coefficient (PCC) for each pair of proteins, resulting in a GCN. Next, we fit a calibration function to convert the PCC scores to posterior probabilities based on KEGG pathways. This pipeline is applied separately to each of the two matrices obtained from the Human Protein Atlas and PRIDE. Finally, we combine the posterior probabilities from the two networks to produce a single, combined network. While the network provided in this paper is based on these specific datasets, the method is applicable to scRNA-seq datasets in general (Figure 2).

Here, we present FAVA, a method to infer Functional Associations using Variational Autoencoders from omics data. We show that i) the method can efficiently handle large-scale scRNA-seq and proteomics data; ii) it outperforms traditional co-expression analysis and the state-of-the-art methods by a considerable margin on both real and simulated data; iii) it can predict high-confidence functional associations, and iv) the resulting networks provide good coverage also for understudied proteins with 4,210 interactions between 1,039 understudied proteins. FAVA is available as a Python package on PyPI, via the command pip install favapy, and it can be used through direct command line or as a module in Python scripts. In addition, the Python module can take as input either a counts/abundance matrix or an AnnData object (24), making it compatible with the most popular single-cell analysis pipelines in Python (e.g. SCANPY (25)).

## Results

### Overview of the FAVA method

FAVA utilizes scRNA-seq data from the Human Protein Atlas (26) and proteomics data from the PRIDE database (27) as inputs to train two separate VAEs (**Figure 1**, top and main box). VAEs are then used to learn the distribution of the input data and create a meaningful low-dimensional representation (i.e. the latent space). Lastly, all-against-all PCCs are calculated in the latent space to quantify similarity in expression between any two genes/proteins. The resulting adjacency matrix is directly translated to a weighted GCN, one for each data type. To combine the two networks, we first convert the Pearson scores into posterior probabilities of belonging to the same biological pathway (KEGG map) by fitting a calibration function (**Figure 1**, equation plot). If an interaction is supported by both scRNA-seq and proteomics data, the two probabilities are combined. The resulting combined network (**Figure 1**, bottom) thus, takes advantage of the complementary nature of the two datasets and is enriched for understudied proteins.

### The FAVA software

The method is implemented in Python and makes use of the Keras deep-learning framework for training VAEs. The software is available via PyPI as ‘favapy’ and is distributed under the MIT open source license. It can be used either directly from the command line, taking an expression matrix file as argument, or as a module in Python scripts. In the Python module, the input can be either a matrix with counts or abundance values from any omics dataset or an AnnData (24) object. The latter is a generic class for handling annotated data matrices making it simple to perform co-expression analysis from scRNA-seq data using popular pipelines.

### FAVA outperforms state-of-the-art methods for GCN construction from individual scRNA-seq datasets

To assess the performance of FAVA on single-cell RNA sequencing (scRNA-seq) data, we tested the method on five diverse datasets: 1,653 (1.5k) human glioblastoma multiforme cells (obtained via 10x Genomics), 2,616 (2.5k) human squamous cell lung carcinoma cells (using Chromium X chemistry), 2,700 (3k) single peripheral blood mononuclear cells (PBMC) (sequenced with the Chromium kit from 10x Genomics), 6,647 (6.5k) human pancreatic tumor cells (isolated with the Chromium Nuclei Isolation Kit), and 30,478 (30k) PBMC (sequenced with the Evercode™ WT kit from Parse Biosciences). For more details on the datasets see Methods. These datasets were selected to cover a range of sizes, origins, and chemistries; moreover, some of them have been used as benchmarks in other studies (25,28). The performance of the resulting gene co-expression networks was evaluated against five gold standard benchmarks, namely manually annotated pathways from KEGG (29) and Reactome (30) physical protein interactions from Complex Portal (31), BioGRID (32) and hu.MAP 2.0 (8). For comparison, we also generated networks using three other GCN methods that are either commonly used or specifically designed for scRNA-seq data, namely PCC, hdWGCNA (13), and scLink (14).

Figure 2 shows that FAVA outperforms the other methods by a wide margin on both datasets and with respect to both gold standards. For a simplified interpretation of the curves, we here focus on the true positive/false positive (TP/FP) ratios in the top 10,000 interactions from each method. On the small dataset, FAVA gave a TP/FP ratio of 3.9 (4,588/1,168) according to the KEGG benchmark (Figure 2a), which is 3.9 times better than the second best method, scLink (1,878/1,860=1.0). For the BioGRID benchmark, the corresponding ratios are 0.29 (2,266/7,734) and 0.1 (1,014/8,986) for FAVA and scLink, respectively (Figure 2b). On the large dataset, which was too large for scLink to process, FAVA also produces the best results with TP/FP ratios of 0.55 (988/1,800) and 0.04 (430/9,570) for the KEGG and BioGRID benchmarks, respectively (Figure 2c**,d**). By comparison, two other methods barely manage to perform better than random.

**Figure 2.**
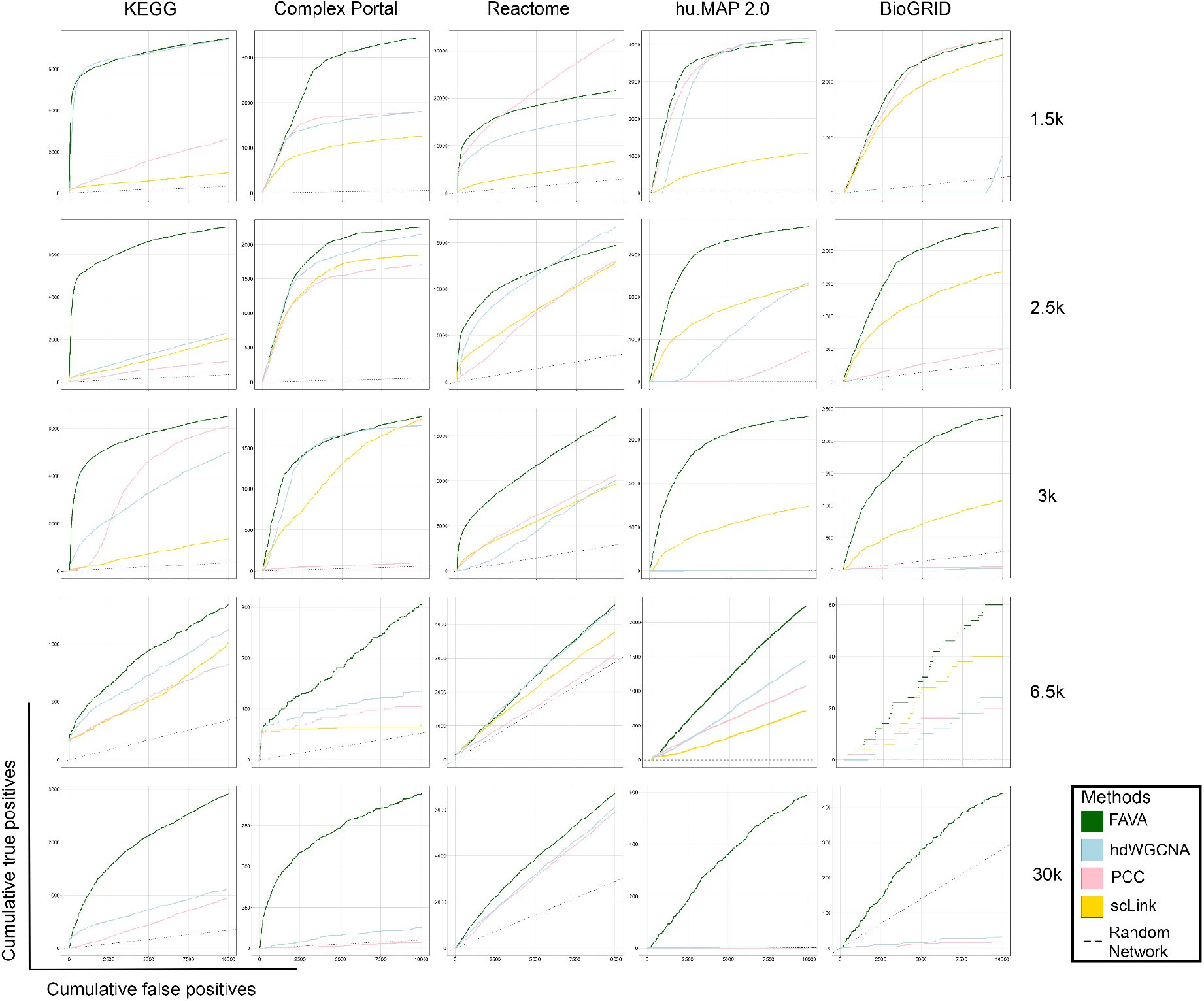
Comparison of the performance of FAVA and the other gene network inference methods on individual scRNA-seq expression data. The plots show method accuracy on GCNs based on different benchmark sets, namely KEGG, Complex Portal, Reactome, hu.MAP 2.0, and BioGRID. Each dataset represents different cell sizes: 1.5k (approximately 1,500 cells of human glioblastoma multiforme obtained through 10x Genomics), 2.5k (approximately 2,500 cells from human squamous cell lung carcinoma using Chromium X chemistry), 3k (2,700 single Peripheral Blood Mononuclear Cells [PBMC] sequenced with the Chromium kit from 10x Genomics), 6.5k (approximately 6,500 cells from human pancreatic tumor isolated with Chromium Nuclei Isolation Kit), and 30k (30,478 PBMC v2 sequenced with the Evercode™ WT kit from Parse Biosciences). As it was not possible to run scLink on the large 30k dataset, this method is only included in the plots for the four smaller datasets. For KEGG and Reactome pathways, a True Positive means that both genes/proteins in a pair are found in the same KEGG/Reactome map, and a False Positive is when both genes/proteins are found in different KEGG or Reactome maps, respectively. For Complex Portal, a True Positive means that both genes/proteins in a pair are found in the same complex, and a False Positive is when both genes/proteins are found in different complexes. For BioGRID and hu.MAP 2.0, a True Positive is a pair of genes/proteins that is also present in the BioGRID/hu.MAP 2.0 interactome, while a False Positive is a pair not present in BioGRID or hu.MAP 2.0, respectively.

We thus conclude that the GCNs produced by FAVA based on scRNA-seq data are better than the corresponding networks from other methods at predicting which proteins function in the same pathway as well as which proteins bind to each other. While neither gold standard is a perfect ground truth for evaluating GCNs, the consistency of the results on two very complementary gold standards and datasets is reassuring.

These findings emphasize two key observations. Firstly, regardless of the dataset size or benchmark sets used, FAVA consistently outperforms other methods. Secondly, the baseline method, PCC, emerges as the second-best performer across various datasets and benchmark sets.

To further investigate the impact of the dataset size on network quality, we subdivided the HPA single-cell atlas into individual tissues and ran FAVA on each. Intriguingly, our findings challenge the notion that a larger number of cells inherently leads to improved performance (Supplementary Figure S1). Specifically, we observed that networks derived from a smaller number of cells exhibited superior performance compared to some others constructed from larger cell populations. This observation underscores the fact that data quality cannot be solely attributed to the quantity of cells, but rather, various dataset-specific characteristics play a substantial role in determining analysis accuracy.

Precision curves, for the pathway databases KEGG and Reactome, have been included in the supplementary material (see Supplementary Figure S2). We did not include hu.MAP 2.0, BioGRID, and Complex Portal in these plots, as they primarily consist of physical interactions, while co-expression networks. Counting all non-physical interactions predicted by FAVA or other co-expression methods as false positives would inherently result in dramatic underestimation of the precision of all methods.

### FAVA performs better on simulated datasets for Gene Regulatory Network predictions

Another common way to assess the performance of computational methods is using simulated data. A few simulators exist that can create simulated expression matrices from Gene Regulatory Networks (GRNs). Even though GRNs remain outside of the scope of methods constructing GCNs, we decided to evaluate their performance on simulated data created with two algorithms, namely scMultiSim (33) SERGIO (34). The results show that FAVA outperforms hdWGCNA, scLink, and PCC for prediction of GRNs from both simulation algorithms (Figure 3). In both cases, FAVA has the best performance with a TP/FP ratio of 0.011 (116/9,884) and 0.02 (208/9,792) for the datasets simulated by scMultisim and SERGIO, respectively. For the scMultisim dataset, scLink gave the second best performance (74/9,926=0.007) (Figure 3a), whereas the ranking of the other methods is less clear on the SERGIO dataset (Figure 3b).

**Figure 3.**
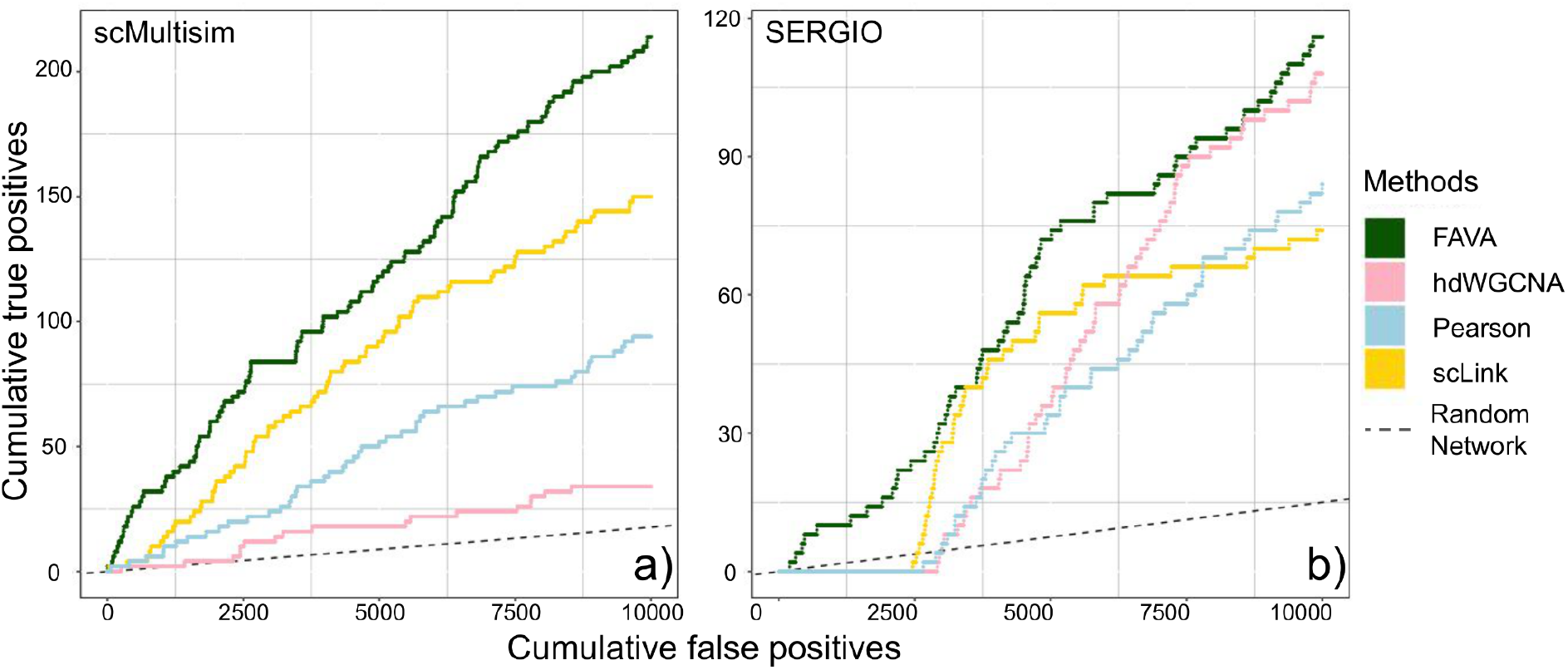
Performance of the methods on simulated data. a) Method performance on simulated data from the scMultisim package. FAVA outperforms the other algorithms with ∼1.6 times better performance than the second best (scLink). b) Methods performance on simulated data from the SERGIO package. FAVA outperforms the other algorithms, whereas the ranking of the other methods is complicated.

We tried to also use a third simulation algorithm, BEELINE (35), which is a well-known framework for benchmarking the performance of computational methods in constructing GRNs. BEELINE contains pre-implemented algorithms dedicated to GRN prediction and suitable expression datasets and benchmarks. Unfortunately, we were not able to make BEELINE run correctly.

In our study, we recognize the fundamental distinction between GRNs and functional association networks, with the former focusing on regulatory interactions among genes and the latter encompassing a broader range of functional associations. Consequently, there is no fair way to compare methods designed to produce these two different types of networks. While benchmarking all available GRN methods on functional association benchmarks is possible, GRN methods should be expected to underperform compared to methods like FAVA, which produce much more inclusive co-expression networks. Conversely, GRN methods ought to outperform FAVA on benchmark sets specifically tailored for regulon analysis.

### Application of FAVA to ∼0.5M single cells and ∼32k proteomics studies

To assess the ability of FAVA to process huge, diverse omics datasets and infer novel functional associations, we applied the method to the single-cell dataset from the Human Protein Atlas, and to proteomics data on human samples from the PRIDE database. We evaluated the quality of the resulting GCNs by benchmarking them against pathways from the KEGG database (Methods).

Here we show that the network derived from scRNA-seq data (Figure 4, green continuous line) captures a very large number of high-confidence interactions in KEGG; it is possible to obtain more than 5,000 TPs with less than 500 FPs, corresponding to a precision of more than 90%. The proteomics-based network (Figure 4, blue continuous line), by contrast, provides fewer high-confidence interactions but more interactions at lower confidence, with the two methods performing on par at ∼3,000 FPs. Comparing the FAVA results to those obtained from PCC on the single-cell and proteomics data (Figure 4, dashed lines), shows that FAVA performs better by a wide margin on both types of data, and especially on scRNA-seq. We did not include scLink and hdWGCNA in this comparison, because the former is unable to handle so large datasets, and both are designed specifically for scRNA-seq data and are thus not applicable to the proteomics dataset.

**Figure 4.**
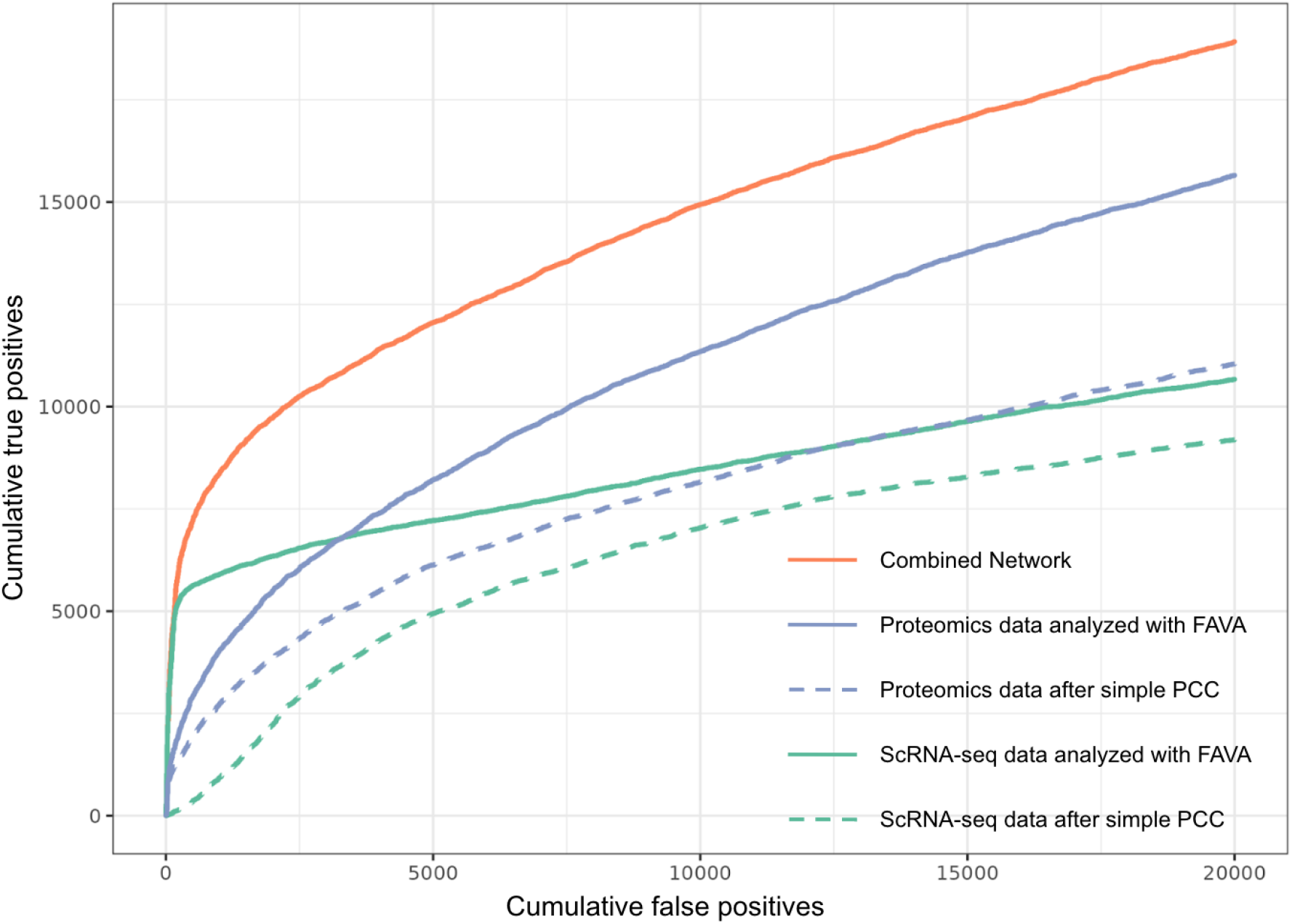
Comparison of FAVA’s results against networks obtained by calculating PCC. The plots show how many known interactions can be predicted according to the KEGG database with the two different methods. True Positive is a pair of proteins when both proteins are found in the same KEGG map. False Positive is a pair of proteins when the two proteins are found in different maps of KEGG. Blue continuous line: Benchmark network from proteomics data after applying FAVA; Blue dashed line: Benchmark network from proteomics data after applying traditional PCC; Green continuous line: Benchmark network from scRNA-seq data after applying FAVA; Green dashed line: Benchmark network from scRNA-seq data after applying traditional PCC; Orange: Resulting network after combining scRNA-seq and proteomics networks coming from FAVA, using KEGG for score calibration (Methods).

### Combined network from scRNA-seq and proteomics data

Given the complementary nature of the networks based on scRNA-seq and proteomics data individually, we decided to combine them into a single network. As the PCC scores from FAVA cannot be assumed to be directly comparable across the two networks, we converted them to probabilistic scores by calibrating both on the KEGG benchmark (Methods). These calibrated scores were then combined to produce a single network based on scRNA-seq as well as proteomics data. Figure 4 shows the performance of the combined network (orange continuous line) derived from both the scRNA-seq and the proteomics networks (green and blue continuous lines, respectively). As should be expected, the combined network shows better agreement with KEGG than either of the individual networks that it was created from. Crucially, we do not consider only the intersection of the networks, which could result in a sparse network with reduced sensitivity (recall); we consider all interactions from both types of data (i.e., the union), combining scores for interactions supported by both types of data.

As was the case for the individual networks, the benchmark against KEGG can also be used to assign probabilistic scores to the interactions in the combined network. At a 15% confidence cutoff (corresponding to the low-confidence cutoff of STRING), the combined network consists of a total of 511,048 associations for 16,790 proteins. The network can be further filtered to obtain higher confidence networks, depending on the concrete use case; in case of proteins with many associations, one will generally want to focus on the highest scoring ones. The combined network has 52,953 associations between 8,191 proteins at the medium confidence cutoff (40%), and even at the high confidence cutoff (70%), it provides 24,182 interactions among 4,166 proteins. The complete combined network, comprising connections with a confidence score of 15% or higher, can be downloaded in its entirety (doi: 10.5281/zenodo.6803472).

### No overfitting on scRNA and proteomics data

In FAVA, separate VAEs are trained to learn the distribution of individual datasets in an unsupervised manner, and the trained models are not applied to unseen data. As no fitting takes place on benchmark datasets, any overfitting of the VAEs to the input data should thus not result in the good performance that we observe. Rather, overfitting should result in a less meaningful latent space and consequently worse performance. To nonetheless evaluate if overfitting happens, we chose a scRNA-seq dataset (36) and randomly split it into a training and a test set, consisting of 67% and 33% of the original dataset, respectively. We trained the VAE only on the training set and tried to predict interactions from both the training and the test set. Similar performance was obtained for both training and test sets (Supplementary figure S3), suggesting that our models do not overfit the data and that they are even able to generalize to unseen data, although such generalization may not be necessary for the specific use cases presented.

### Associations for understudied proteins

Given the nature of the data that the combined network is based on, it should be free of the inherent literature bias, which many other protein networks suffer from. Additionally, the network incorporates numerous high-confidence interactions, rendering it highly suitable for the exploration and understanding of the functions associated with understudied proteins.

To explore this, we use the definition of “understudied proteins” from the National Institutes of Health (NIH) funded consortium “Illuminating the Druggable Genome” (IDG) (37). IDG provides information on human proteins (targets), dividing them into four levels for drug development: “Tclin” are drug targets with known mechanisms of action approved by DrugCentral; “Tchem” are targets selected from ChEMBL/DrugCentral or manually curated from other sources; “Tbio” are biologically characterized targets without known drug or small molecule activities that meet specified activity thresholds; and “Tdark” are targets with little known information and not meeting the drug/small molecule activity criteria. In our analysis we used the list of 6,000 proteins that have been designated as Tdark (“dark” targets) by IDG (3). Looking these up in the combined FAVA network, based on both scRNA-seq and proteomics data, revealed 4,210 predicted interactions between 1,039 of the understudied proteins (Figure 5, purple nodes) and 611 other, better studied proteins (Tclin) with at least 70% confidence. Next, we compared this network to hu.MAP 2.0 (8) to evaluate both the agreement and the complementarity of the two networks. Since hu.MAP 2.0 is a physical interaction network, it should contain only a subset of the interactions from the combined FAVA network. In the Supplementary figure S4 we report the precision and recall calculations of our network on hu.MAP 2.0 edges, showing the agreement between the two networks. Furthermore, comparing the two networks, both at 70% confidence, we found that they agree on 1,284 edges (Figure 5, dark gray edges) for 520 nodes, which confirms the high quality of both networks and shows that the FAVA network includes physical interactions among the functional associations. Focusing on the understudied proteins, 49 of the 1,039 that are found in the FAVA network are also among the 648 that hu.MAP 2.0 provides physical interactions for (see Supplementary Figure S5). This shows that while there is good agreement between the two networks, they are also highly complementary, illustrating that FAVA provides much needed additional information about different understudied proteins.

**Figure 5.**
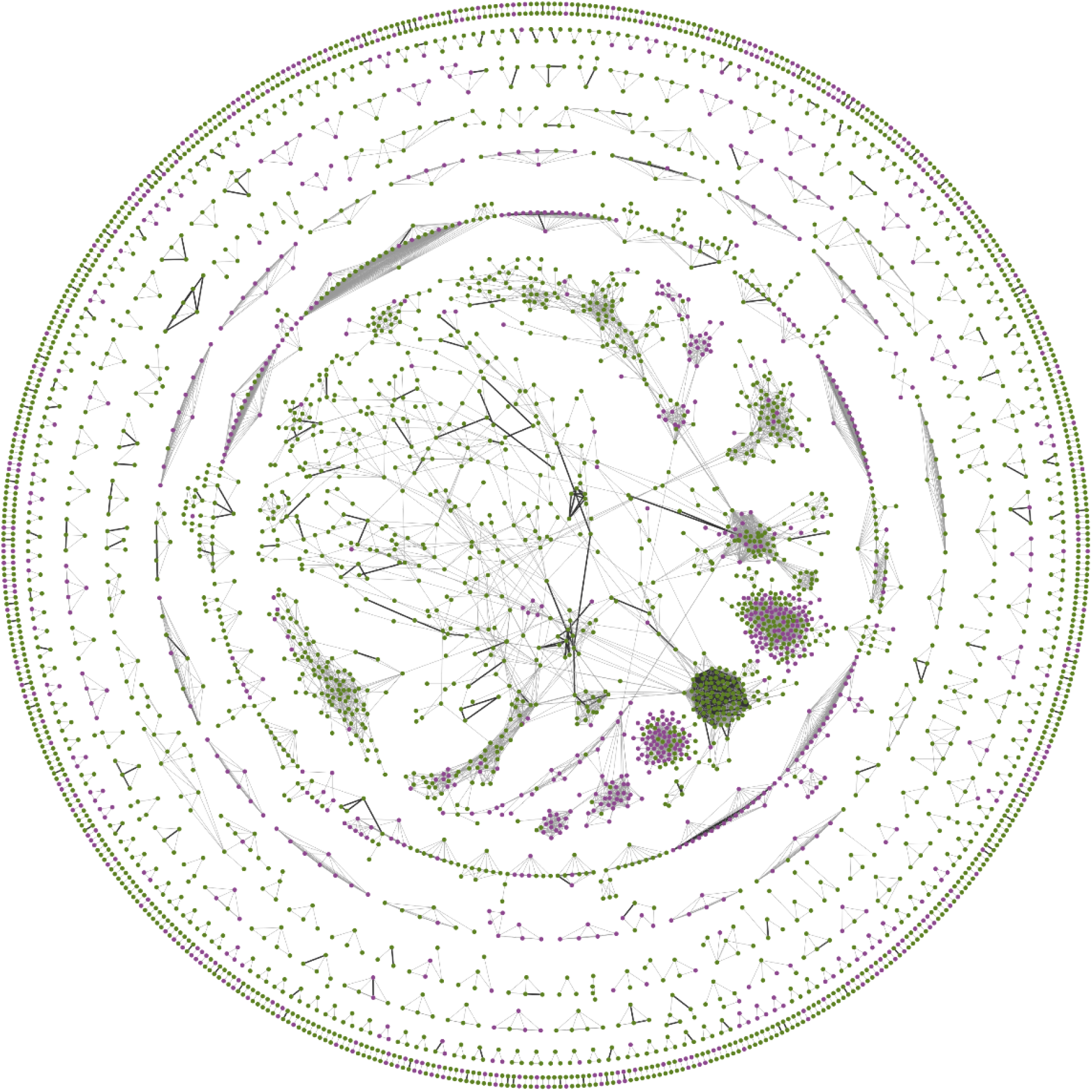
Understudied proteins in the combined FAVA Network. The figure shows all functional associations with a confidence score of at least 70%, based on FAVA analysis of both the scRNA-seq and proteomics data. Understudied proteins according to IDG are highlighted in purple. The edges in the network which are physical interactions according to hu.MAP 2.0 are highlighted with darker, thicker edges. The visualization of the network was made in Cytoscape (38) and the analysis for the IDG target classification was retrieved using Cytoscape stringApp (39).

To evaluate the novelty of the interactions discovered by FAVA, we checked how many of the FAVA edges can be found in the STRING v11.5 and hu.MAP 2.0 databases. Using multiple score cutoffs for STRING, we analyzed the overlapping edges and found that the FAVA network contains 2,747 high-scoring interactions for understudied proteins (Tdark) that are not present in the STRING network at all, and only two of these are found in hu.MAP 2.0 (Supplementary Figure **S6)**. This highlights the value of incorporating co-expression analysis and demonstrates the ability of FAVA to uncover novel interactions, complementing existing experimental and literature knowledge.

### Clustering and functional enrichment analysis

To assess whether functional modules can be detected in the FAVA network, we first clustered the network using the MCL algorithm (inflation value 1.5) (40). This resulted in a network with 341 clusters, 130 of which comprised five or more nodes, and were further analyzed. Next, we used Cytoscape stringApp 2.0 (41) to perform enrichment analysis on the clustered network.

The full enrichment results are available in Supplementary Table 1, Sheet 1. To functionally characterize the clusters, we manually selected highly significant terms from the following seven categories: namely Pathways, UniProt Keywords, Cellular Compartments, Tissues, Biological Processes, Molecular Functions and Diseases. Five selected clusters are shown in Figure 6, with colored rings around the nodes to indicate the most descriptive functionally enriched terms for each cluster. Results for all clusters can be found in Supplementary Table 1, Sheet 2.

**Figure 6.**
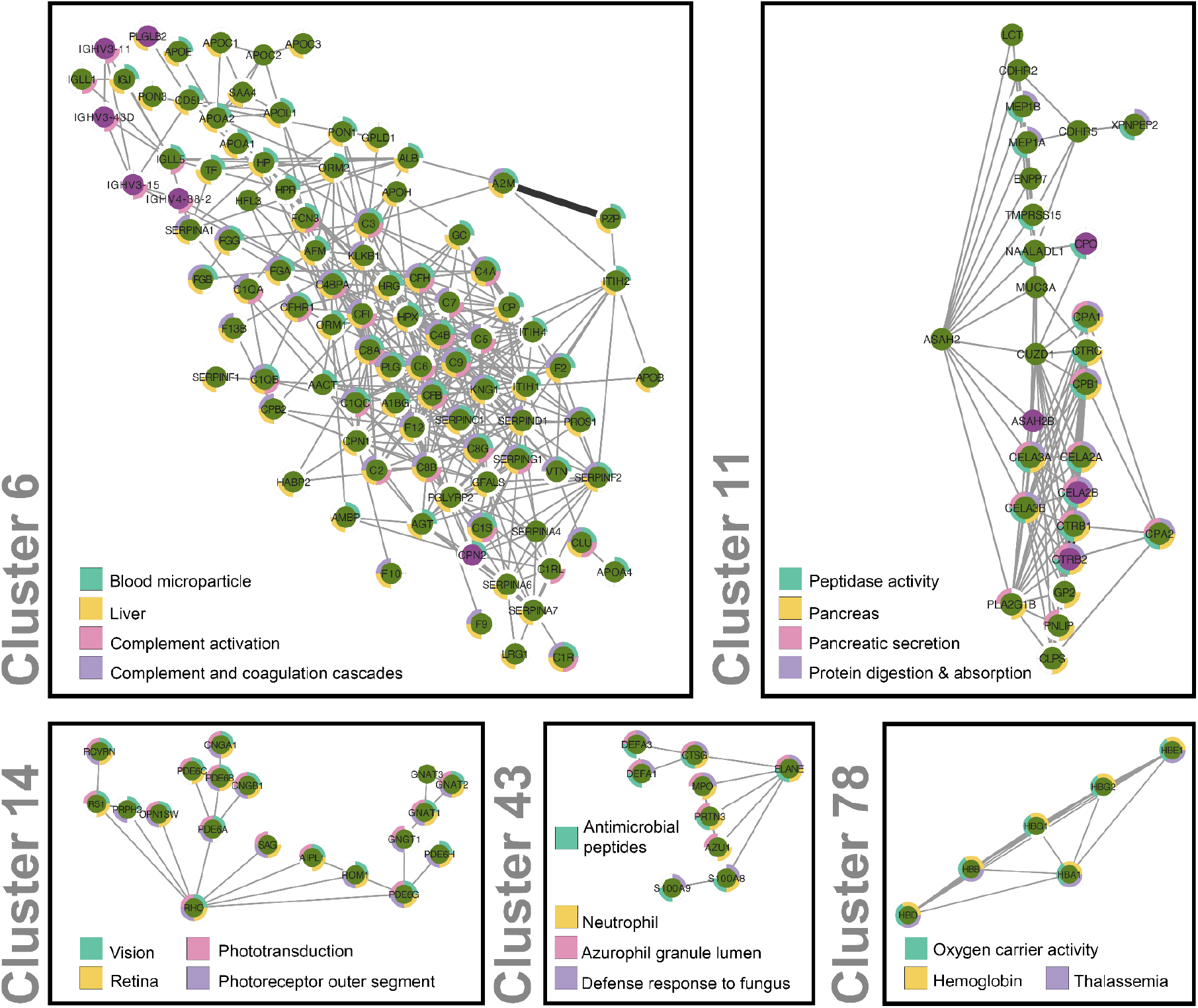
Clustering and functional enrichment analysis of the combined FAVA network. Five selected clusters with five or more nodes, derived utilizing the MCL algorithm were visualized using Cytoscape (38). The visual legend summarizes the enriched terms, depicted as quarters of colored rings around the protein nodes of each cluster. The clusters represent different functions and pathways Clusters 6 and 11 also illustrate how FAVA is able to link understudied proteins (Tdark; purple) to well-annotated ones (green).

## Discussion

Here we introduced FAVA, a new method to build functional association networks from omics data using VAEs. FAVA takes an expression matrix from, for example, scRNA-seq or proteomics experiments as input and uses VAEs to create a low-dimensional representation of it, known as the latent space. It then produces a weighted GCN by calculating PCCs in the latent space. To evaluate the ability of FAVA to handle sparse and redundant data, we have tested it on two scRNA-seq datasets of very different size and compared its performance to that of tools designed or commonly used for the same purpose, namely scLink, hdWGCNA and PCC. We evaluated the methods on two complementary benchmark sets, namely pathways from KEGG and physical protein interactions from BioGRID. Overall, FAVA demonstrated the best performance.

FAVA is an efficient, unsupervised, stand-alone method, which does not rely on the results from other approaches, as the hdWGCNA does. Additionally, it considers the expression count of all genes, while scLink only takes into consideration the expression counts of certain highly expressed genes. With that, FAVA avoids the accumulation of “rich-get-richer” genes/proteins and produces networks that contain both studied and understudied genes/proteins, which is a major issue in the field. Compared to PCC, FAVA is at least nine times faster (1 day ≃ 9 days) and is better at accurately identifying established patterns in the data.

The efficiency of FAVA allows the method to be applied not only to individual studies, but also to large expression atlases. This versatility of FAVA allows the construction of GCNs from vast amounts of data of different types. As illustrated in Figure 5, FAVA generates GCNs from scRNA-seq and proteomics data and outperforms conventional co-expression methods. The combination of these networks offers new perspectives on uncharacterized proteins. This highlights the ability of the method to handle data beyond scRNA-seq. Overall, the efficiency of FAVA makes it a powerful and flexible approach for gene expression studies.

FAVA also achieves higher accuracy in gene network construction on simulated datasets, as presented on Figure 3. However, we remain skeptical about these results, because the simulation algorithms used were not designed for the same purpose as the methods discussed. Indeed, we utilized simulation algorithms that create expression datasets based on GRNs. Then we evaluated the performances of FAVA, scLink, hdWGCNA and PCC, methods developed for building GCNs, regarding their accuracy on GRN predictions. We concluded that while none of the methods had accuracy suitable for this task, FAVA performed better than the other methods.

## Conclusions

In this work, we have shown that variational autoencoders can be used to model single-cell RNA-seq and proteomics data for prediction of functional associations. We have applied this method, which we call FAVA, to large compendiums of scRNA-seq data as well as proteomics data. The results show that FAVA scales to such large datasets, and that the resulting networks are considerably more reliable than those obtained from traditional co-expression methods to the same data. We have moreover shown that combining FAVA results from both data types provides an even more comprehensive network, and that this network can be used to associate understudied proteins with better studied ones, thereby providing hints to their possible functions. We make these networks publicly available along with the Python implementation of FAVA, which can be installed as the PyPI package ‘favapy’.

## Methods

### Datasets

#### Human Glioblastoma Multiforme (∼1,500 cells)

This dataset consists of 2,000 sorted cells from Human Glioblastoma Multiforme, prepared using the Chromium Next GEM Single Cell 3ʹ Reagent Kits v3.1. The cells were obtained from a 71-year-old male donor and provided by 10x Genomics through Discovery Life Sciences. The libraries were sequenced on an Illumina NovaSeq 6000, with an average sequencing depth of 72,259 reads per cell. The analysis was performed using Cell Ranger 6.0.0 with the parameter “--expect-cells=2000”. After filtering the number of cells decreased to 1,653. https://www.10xgenomics.com/resources/datasets/2-k-sorted-cells-from-human-glioblastoma-multiforme-3-v-3-1-3-1-standard-6-0-0

#### Human Squamous Cell Lung Carcinoma (∼2,500 cells)

This dataset is part of the Alternative transcript isoform detection with single cell and spatial resolution. It includes cryopreserved, dissociated tumor cells from Stage III squamous cell lung carcinoma (lung cancer DTCs) obtained from Discovery Life Sciences. The dataset consists of 5’ Single Cell Gene Expression Libraries generated from approximately 4,000 cells, with 2,616 cells successfully recovered. The libraries were prepared following the Chromium Single Cell 5’ Reagent Kits User Guide (v2 Chemistry Dual Index) and sequenced on an Illumina NovaSeq 6000, achieving an average read depth of around 25,000 reads per cell. https://www.10xgenomics.com/resources/datasets/3k-human-squamous-cell-lung-carcinoma-dtcs-chromium-x-2-standard

#### Small PBMCs dataset (3,000 cells)

This dataset consists of 2,700 single Peripheral Blood Mononuclear Cells (PBMC) that were sequenced using Chromium kit from 10x Genomics. We downloaded the counts matrix for this freely available dataset from 10X Genomics: https://cf.10xgenomics.com/samples/cell/pbmc3k/pbmc3k_filtered_gene_bc_matrices.tar.gz

#### Human Pancreatic Tumor (∼6,500 cells)

This dataset comprises 5,000 human pancreatic tumor cells isolated using the Chromium Nuclei Isolation Kit. The tumor tissue, obtained from Discovery Life Sciences, was processed following the Chromium Nuclei Isolation Reagent Kits Sample Prep User Guide. Gene expression libraries were prepared using the Chromium Single Cell 5’v2 Reagent Kits and sequenced on an Illumina NovaSeq instrument with a target of 20,000 reads per cell. The data were processed using Cell Ranger 7.0.0, resulting in the detection of 6,647 cells, with a median of 1,776 genes and 3,465 UMIs per cell. The sequencing depth averaged 56,169 reads per cell in a paired-end, dual indexing configuration. https://www.10xgenomics.com/resources/datasets/5k-human-pancreatic-tumor-isolated-with-chromium-nuclei-isolation-kit-3-1-standard

#### Large PBMCs dataset (∼30,000 cells)

This larger dataset of 30,478 Peripheral Blood Mononuclear Cells (PBMC v2) was sequenced after applying the Evercode™ WT kit. We downloaded the counts matrix for this freely available dataset from Parse Biosciences: www.parsebiosciences.com/datasets

#### Human Protein Atlas – Single-cell RNA-seq read-count data

We obtained the single-cell dataset from the Human Protein Atlas (https://www.proteinatlas.org/humanproteome/single+cell+type), a public resource that provides transcriptomics and spatial antibody-based proteomics profiling of human tissues. Their single-cell transcriptomics atlas combines data from 26 datasets. The matrix gives read counts for 19,670 human protein-coding genes in 566,109 individual cells grouped into 192 cell type clusters. Information about the external datasets and processing of the data is described in detail in (42).

#### PRIDE EMBL-EBI – Proteomics dataset

We obtained our proteomics dataset from The PRoteomics IDEntifications (PRIDE - https://www.ebi.ac.uk/pride/) database, the world’s largest data repository of mass spectrometry-based proteomics data (27). Specifically, we used 633 human proteomics project experiments with a total of 32,546 runs and reanalyzed them using ionbot (43) with an FDR threshold of 0.01 (44), resulting in a total of 154,885,151 peptide spectrum matches for 18,846 proteins. For the full list of projects, runs, and general statistics of the search see supplementary material in Zenodo (doi: 10.5281/zenodo.6798182).

#### Simulated Datasets

To benchmark the GCN methods on simulated data, we used two recent simulators, SERGIO and scMultiSim, to create single-cell expression matrices based on GRNs. We then used FAVA, scLink, hdWGCNA, and PCC to create GCNs, and benchmarked these on the ground-truth GRNs.

#### scMultisim

To construct an expression matrix with scMultisim, we follow the pipeline described here: https://zhanglabgt.github.io/scMultiSim/vignettes/sim_new.nb.html. To allow for generation of a matrix with 2,000 cells and 1,000 genes we ran the following command: `options_ = list(rand.seed=0, GRN=GRN_params_100, num.cells=2000, diff.cif.fraction=0.8, intrinsic.noise=1).` The given GRN contains 1,432 interactions.

#### SERGIO

From SERGIO, we used the `SERGIO_noised_1200G_9T_300cPerT_dynamics_6_DS83` file, which is a simulated matrix populated with 300 cells and 1,200 genes and includes technical noise. The package also offers the equivalent GRN with 2,713 interactions, from which the simulated matrix is derived.

#### FAVA method

We have developed a novel method for construction of GCNs from huge omics datasets with high data sparseness and redundancy. The method makes use of VAEs to perform dimensionality reduction and subsequently scores co-expression in the latent space.

#### Pre-processing count data

The first step is to *log_2_* normalize the count matrices. By log-transforming the values, we model proportional changes rather than additive changes and we help the model focus on the biologically relevant differences rather than the extreme values. The general recommendation is to ensure that the data lies in the range of the function we are using to approximate it, in our case a neural network with sigmoid activation function on the output layer. Then, we normalized the values to the [0,1] range by dividing by the max value of the row that each value belongs.

#### Dimensionality reduction using Variational Autoencoders and Hyperparameters

To compress the high-dimensional expression matrices into lower-dimensional latent spaces, we make use of Variational Autoencoders (VAEs). Using VAEs, the latent representations are regularized by minimizing the regularization loss, implemented with the Kullback–Leibler (KL) divergence between the latent and a prior, in this case a normal distribution with mean 1 and 0.1 standard deviation. The VAE loss is composed by the reconstruction and regularization losses weighted sum, where the weights are set to 0.9 and 0.1 respectively, obtained through hyperparameter tuning. Regarding the VAEs architecture, VAE performance was evaluated with different layers. We use a single hidden layer in FAVA. In all cases, we used the Rectified Linear Unit (ReLU) function (45) for all layers, except for the sigma and mu encoding layers and the last layer in the decoder. In the output layer, considering the 0-1 scaling of the inputs, a sigmoid function was used to generate values between 0 and 1, whereas linear activation was set for the mu and sigma layers. To train the VAE, we used the Adam optimizer with a learning rate of 10^−3^. The VAE model was implemented in Keras (https://keras.io/).

#### Pearson Correlation Coefficient pairwise on the latent space

Having produced a regularized latent space that follows Gaussian distribution from the VAE, we calculate all pairwise PCCs between proteins in the latent space. That outputs a list of protein pairs with an assigned score, showing the proximity of the two proteins in the latent space. Based on this score, we sort all protein pairs and create a ranked list, in which numbers closer to 1 represent higher proximity in the latent space and thus, expression similarity. Finally, we benchmark the resulting ranked list against the KEGG database (29) to quantify how well the predicted interactions agree with what is known.

#### Co-expression scoring in the latent space

Having produced a regularized latent space that follows Gaussian distribution from the VAE, we calculate all pairwise PCCs between proteins in the latent space. That outputs a list of protein pairs with an assigned score, showing the similarity of the two proteins in the latent space. Based on this score, we sort all protein pairs and create a ranked list, in which numbers closer to 1 represent higher similarity in the latent space and thus, expression similarity.

#### Generalization of FAVA algorithm on unseen data

To show that the FAVA method can even generalize to genes not included in training the VAEs we employed another dataset from GEO (GSE75748 (36)). We split it into a 67% train set and a 33% test set. We, then, trained our VAEs on the train set and we separately applied the trained model to both the train and test sets. Two different networks were generated by the two different latent spaces that were created. The accuracy of these networks was evaluated using KEGG (see Supplementary figure **S3**).

#### Network construction with FAVA

FAVA is available as a PyPI package and can be used either in the terminal or as a module in a Python script. In both datasets, we applied FAVA from terminal with the following command: `favapy <path-to-data-file> <path-to-save-output>`, as described here: https://pypi.org/project/favapy/

### Other methods and networks

#### scLink

To create the network of the small dataset using the scLink method, we followed the pipeline described here: https://cran.r-project.org/web/packages/scLink/scLink.pdf. Thus, we normalized the data with the package’s function and filtered for the 500 most highly expressed genes, as recommended by the authors: `sclink_norm(count, scale.factor=1e+06, filter.genes = TRUE, n = 500)`. Then we calculated the correlation matrix with the function: `sclink_cor(expr = count.norm)`. We were not able to process the large dataset on a server with 180GB RAM.

#### hdWGCNA

To create the networks using hdWGCNA, we first had to analyze the data using the Seurat pipeline. Thus, we followed the tutorial provided by Seurat here: https://satijalab.org/seurat/articles/pbmc3k_tutorial.html. That creates a Seurat object (seurat_obj) which can be utilized further by hdWGCNA. Afterwards, we followed the recommendations of the authors to construct the network as described in their tutorial here: https://smorabit.github.io/hdWGCNA/articles/basic_tutorial.html. Therefore, we ran in R the following commands: seurat_obj = SetupForWGCNA(seurat_obj, gene_select = “fraction”, fraction = 0.05) seurat_obj = SetDatExpr(seurat_obj, assay = ‘RNA’, slot = ‘data’) seurat_obj = TestSoftPowers(seurat_obj, networkType = ‘unsigned’) Based on the results coming from the `GetPowerTable(seurat_obj)` function, we selected the soft_power parameter (`4` for the small dataset and `7` for the large dataset). Final step was to construct the network: `seurat_obj = ConstructNetwork(seurat_obj, soft_power = soft_power, setDatExpr = FALSE, overwrite_tom = TRUE, networkType = ‘unsigned’)`

#### Pearson correlation coefficient (PCC)

We calculated all-against-all PCC on the same log-transformed counts matrices used as input for FAVA. This was done in order to have a fair comparison with FAVA.

### Benchmark of functional associations

#### KEGG

We benchmark the resulting ranked list against the KEGG database, identical to how functional associations are benchmarked in the STRING database (4), to quantify how well the predicted interactions agree with what is known. To do this, we first map the protein pairs from the methods to KEGG maps. If a KEGG map exists, which contains both proteins of a pair, the pair is counted as a true positive (TP). If both proteins can be mapped to KEGG, but there is no map containing both, the pair is counted as a false positive (FP). Pairs for which one or both proteins cannot be mapped to KEGG are disregarded for benchmarking purposes. Having defined which protein pairs are considered TPs and FPs, we plot the cumulative TP count as a function of the cumulative FP count for the sorted lists of pairs. Furthermore, we generate a plot that illustrates the precision in relation to the TP predictions for the sorted list of protein pairs obtained from the methods.

#### Reactome

Similar to the approach used in benchmarking with the KEGG database, Reactome pathways can be utilized to assess the concordance between predicted interactions and known functional associations. Protein pairs obtained from the prediction methods are mapped to the Reactome pathways. If a Reactome pathway contains both proteins of a pair, it is considered a true positive (TP). However, if both proteins can be mapped to Reactome but no pathway encompasses both, the pair is counted as a false positive (FP). Pairs where one or both proteins cannot be mapped to Reactome are excluded from the benchmarking analysis.

#### Complex Portal

We evaluate the performance of the ranked list by comparing it to the Complex Portal database. Protein pairs obtained from the methods are mapped to the entries in the Complex Portal. If a Complex Portal entry includes both proteins of a pair, it is considered a true positive (TP). If both proteins can be mapped to the Complex Portal but no entry contains both, it is counted as a false positive (FP). Pairs with unmappable proteins are excluded. The evaluation is visualized through a plot showing the cumulative TP count against the cumulative FP count for the sorted lists of pairs. By benchmarking against the Complex Portal database, we assess how well the predicted interactions align with known information regarding physical protein complexes.

#### BioGRID

To measure how accurately the networks from the different methods correlate with BioGRID’s experimental results, we considered the interaction network from BioGRID as our ground truth. If a predicted pair existed in BioGRID, it was labeled as a True Positive (TP). Conversely, if it was not present in BioGRID, it was labeled as a False Positive (FP). The interaction network of BioGRID can be found here: https://downloads.thebiogrid.org/BioGRID/Release-Archive/BIOGRID-4.4.217/

#### hu.MAP 2.0

To evaluate the accuracy of the networks generated by different methods in correlation with experimental results, we utilized the hu.MAP 2.0 database as our benchmark. The hu.MAP 2.0 database provides a comprehensive resource for studying molecular interactions. In this evaluation, we considered the interaction network within hu.MAP 2.0 as our reference or ground truth. For each predicted protein pair, we checked if the pair existed in the hu.MAP 2.0 database. If a predicted pair was present in hu.MAP 2.0, it was labeled as a True Positive (TP), indicating a correct prediction. On the other hand, if a predicted pair was not found in hu.MAP 2.0, it was labeled as a False Positive (FP).

### Score calibration and combination of FAVA networks from atlases

To combine the two networks, we decided to apply the pipeline used in the STRING database. Thus, we first convert the PCC scores from FAVA from each dataset into posterior probabilities of being on the same KEGG map given the PCC (Figure 1). We do this based on the benchmark results described above, by first plotting the local precision within a sliding window (*y = TP/(TP + FP)*) as function the average PCC within the window (*x*). We do this separately for scRNA-seq and proteomics data, and fit the following calibration function to each by minimizing the squared error using simplex optimization: *y* = *a*0 + *a*1 * *x* + *a*2/(1 + *exp*(*a*3 * (*x* − *a*4))), where *a*0 through *a*4 are the parameters that are optimized to fit the points in the plot. Once fitted, we use the resulting calibration curves to convert all PCCs from each dataset to probabilities. When a pair is supported by both scRNA-seq and proteomics data, the two probabilities are combined.

## Availability of data and materials

### Data availability

#### Combined network

The Network: https://doi.org/10.5281/zenodo.6803472

#### The PRIDE database analysis

The full list of projects, runs, and general statistics about the analysis of the data in the PRIDE database: https://doi.org/10.5281/zenodo.6798182

#### Code availability

https://github.com/mikelkou/fava

PyPI: <pip install favapy>

We have created a Jupyter notebook that provides users with a comprehensive analysis toolkit, to facilitate effective interpretation of their networks. This notebook enables users to extract meaningful results from their networks by selecting the desired number of interactions and conducting analyses tailored to their specific context. To access the notebook and learn how to utilize FAVA effectively, please refer to the following link on GitHub: <u>GitHub - How to use FAVA in a notebook</u>

In this notebook we demonstrate also how FAVA can be used via the scverse ecosystem (46), by giving to favapy an AnnData object and creating directly from this a network.

## Supporting information

Protein networks are commonly used for understanding how proteins interact. However, they are typically biased by data availability, favoring well-stu

## Acknowledgments

The authors acknowledge Ralf Gabriels for input on the implementation of the Python package ‘favapy’ and Enrico Massignani for the PRIDE data retrieval.

## Funding

This work was supported by the Novo Nordisk Foundation [NNF14CC0001], [NNF20SA0035590] and EMBO Scientific Exchange Grant 9404 [STF_9404]. R.B. acknowledges funding from the Vlaams Agentschap Innoveren en Ondernemen under project number HBC.2020.2205. Research Foundation - Flanders (FWO) G028821N to L.M., by the European Union’s Horizon 2020 Program (H2020-INFRAIA-2018-1) [823839 to L.M.], and Ghent University Concerted Research Action [grant number BOF21-GOA-033 to L.M.]. Funding for open access charge: Novo Nordisk Foundation [NNF14CC0001] and [NNF20SA0035590]. K.N. has received funding from the European Union’s Horizon 2020 research and innovation program under the Marie Sklodowska-Curie [grant number 101023676].

## Author contributions

M.K. and L.J.J. conceptualized this study. The manuscript was written by M.K. and L.J.J. with assistance and approval from all authors. M.K. developed the favapy Python package. M.K. and P.P.L. designed the architecture of the VAEs. S.R. gave feedback on the VAEs architecture. M.K. collected, processed, and performed network analysis on publicly available datasets. M.K. implemented, processed and performed network analysis on simulated datasets. K.N. performed the clustering and enrichment analysis of the combined network of FAVA. R.B. and L.M. analyzed and provided the proteomics datasets from PRIDE. M.K. collected, analyzed and constructed networks from Human Protein Atlas (single-cells) and PRIDE (proteomics) data.

## Corresponding authors

Correspondence to Lars Juhl Jensen.

## Ethics declarations

### Conflict of interest

None declared.

### Ethics approval and consent to participate

Ethics approval was not applicable for this study.

## Supplementary material

**S1.**
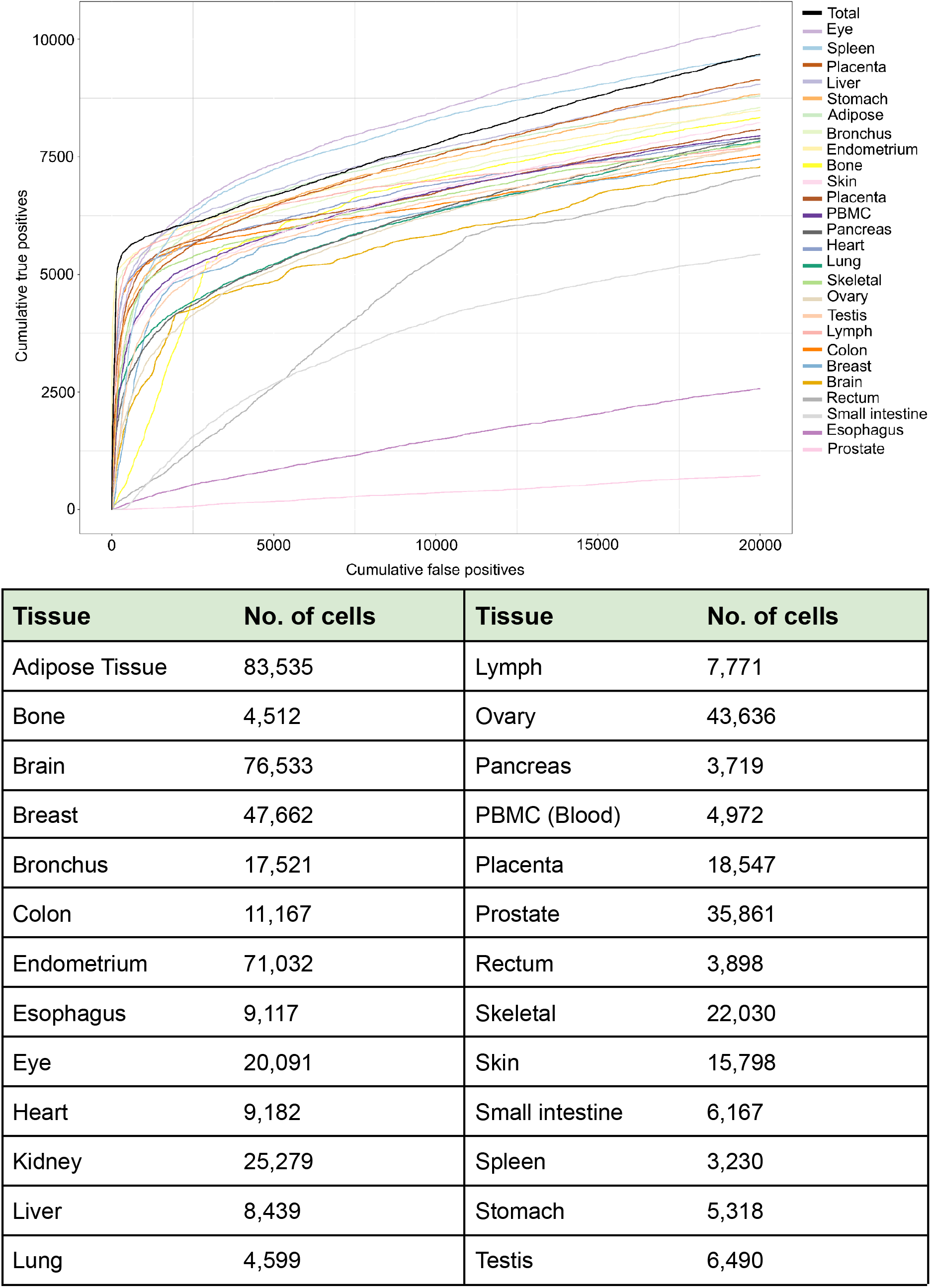
Performance of FAVA Networks on KEGG Pathways: Influence of Cell Numbers on Network Quality. In our benchmarking analysis, we evaluated the performance of FAVA networks on KEGG pathways and observed that the majority of tissues yielded high-quality networks. Surprisingly, we found that the quality of the networks did not straightforwardly correlate with the number of cells used. For instance, the spleen network demonstrated excellent performance despite being constructed from just 3,230 cells. This observation suggests that while having a larger number of cells can be advantageous, it does not necessarily guarantee superior network quality. The fact that the spleen network, with its limited cell count, produced one of the better networks underscores the complexity of the relationship between the number of cells and network performance in FAVA. These findings highlight the need for a nuanced understanding of the factors influencing network quality and support the notion that FAVA can generate robust networks even with modest cell numbers.

**S2.**
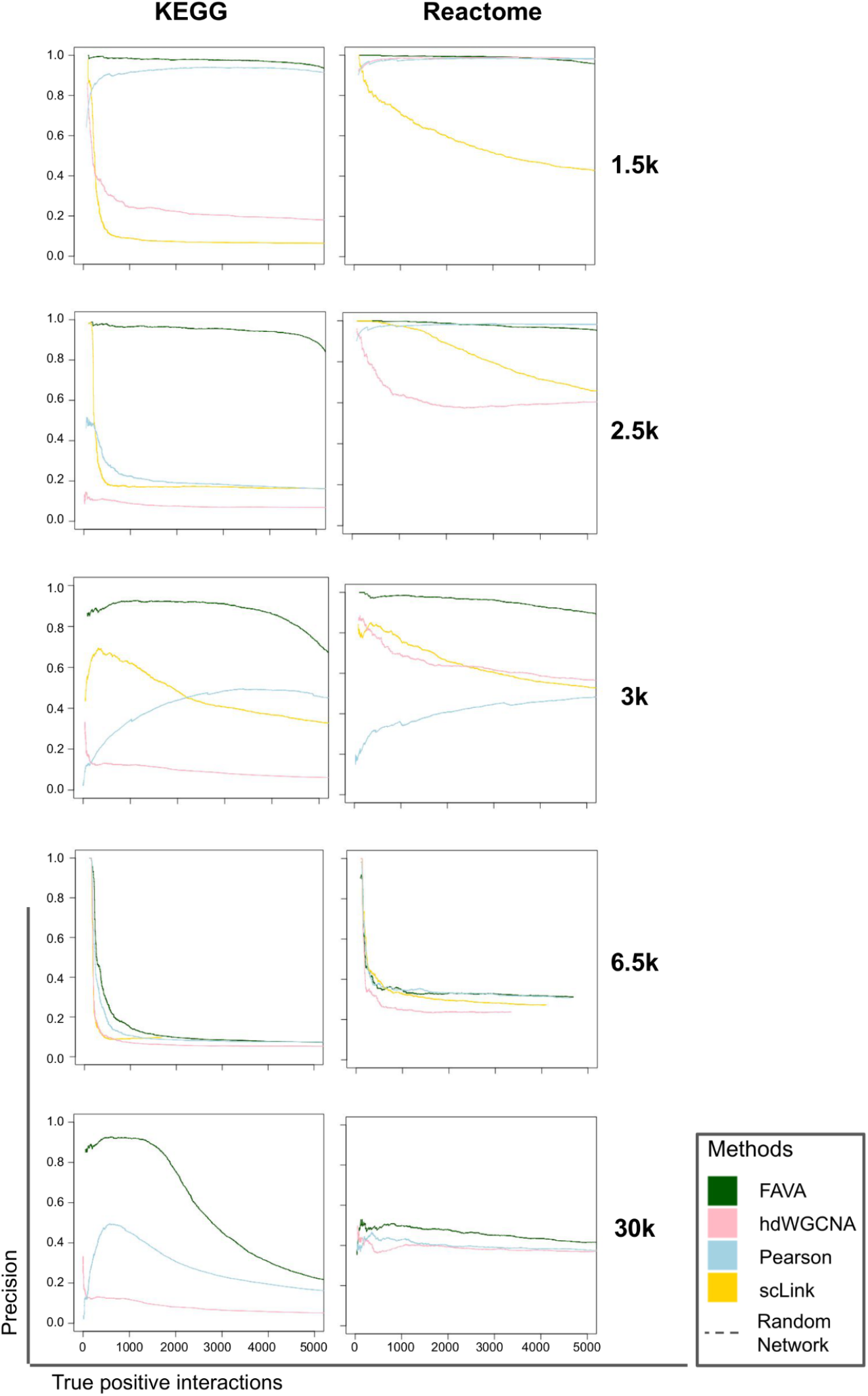
Precision curves. The precision plots for the KEGG and Reactome databases are presented. These plots illustrate the precision values obtained for different methods in relation to the benchmark datasets. The precision values reflect the accuracy of the methods in identifying interactions within the KEGG and Reactome pathways. It is worth noting that the precision curves provided are not precision–recall curves due to challenges in calculating recall based on pathways. The imbalance in the datasets is acknowledged as a concern for ROC curves, which is why the true positives and false positives are explicitly shown in the plots instead of normalizing the data.

**S3.**
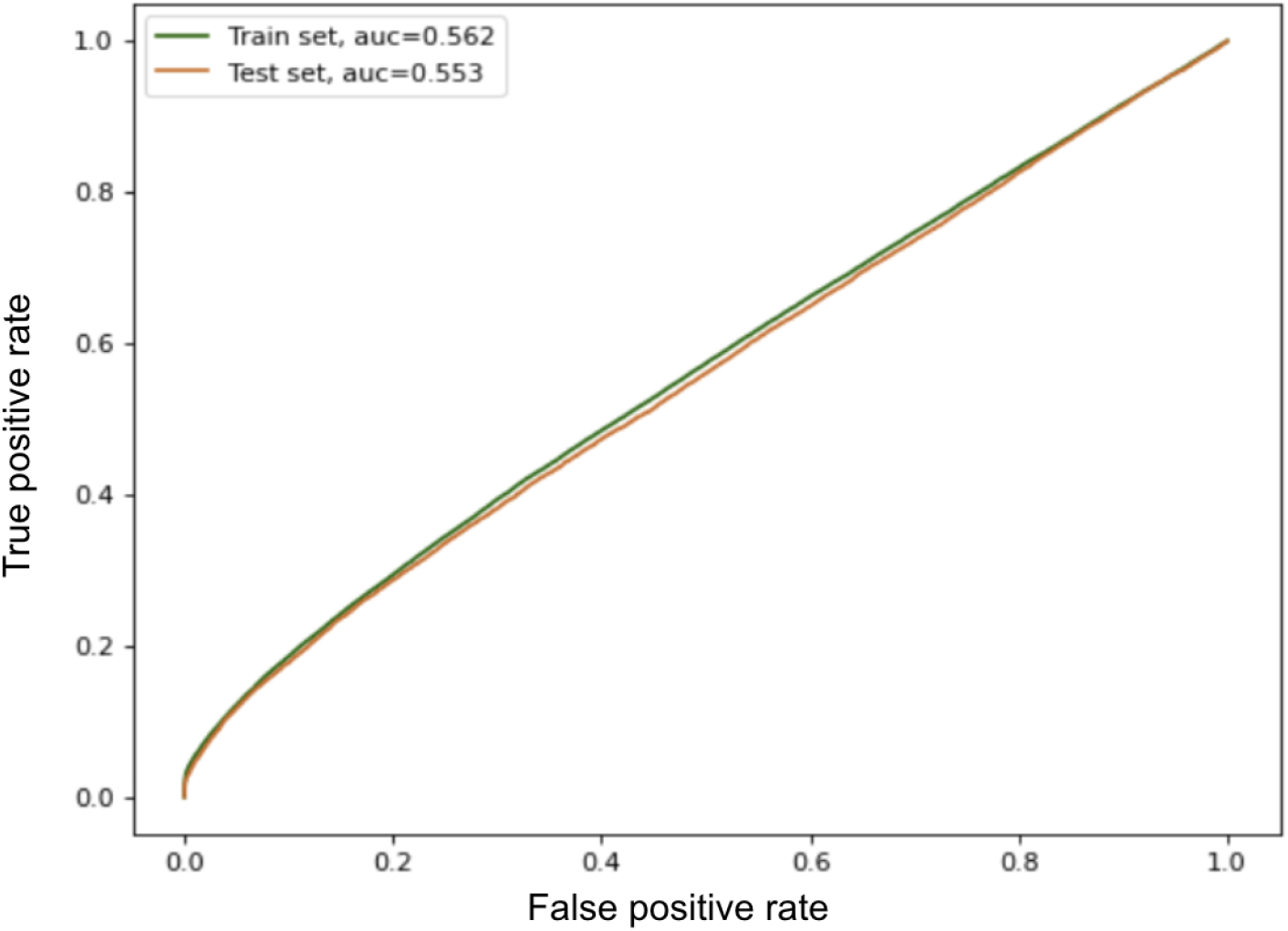
Generalization on unseen data. We split the scRNA-seq dataset (GSE75748) into a training (67%) and a test (33%) set. After training FAVA, the algorithm was applied on both the training and the test set to predict associations. The performance of the predictions is evaluated on KEGG (Methods). The plot shows that our framework is able to generalize when applied on unseen data (i.e. the test set). This is evident from the similarity between the green line (train set) and brown line (test set) in terms of true positive predictions. It is clear that even in the unseen 33% of the dataset, the VAE is able to capture patterns and that the latent representation can be used to predict existing interactions.

**S4.**
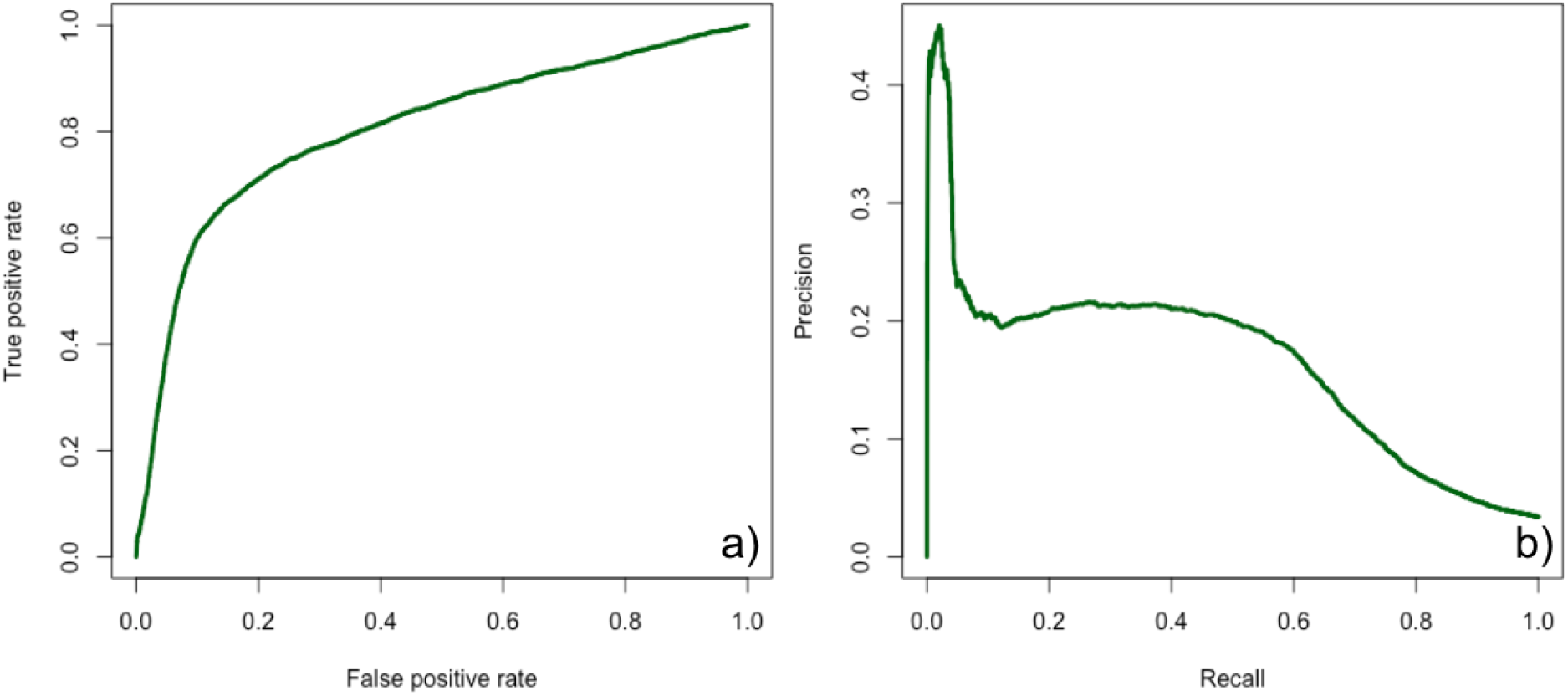
ROC curve, Precision and recall of FAVA’s combined network based on hu.MAP 2.0. We conducted a comparison between our network from single-cells and proteomics data and the hu.MAP 2.0 physical interaction network. We report the a) roc curve and the b) precision and recall calculations of our network on hu.MAP 2.0 edges, showing the agreement between the two networks.

**S5.**
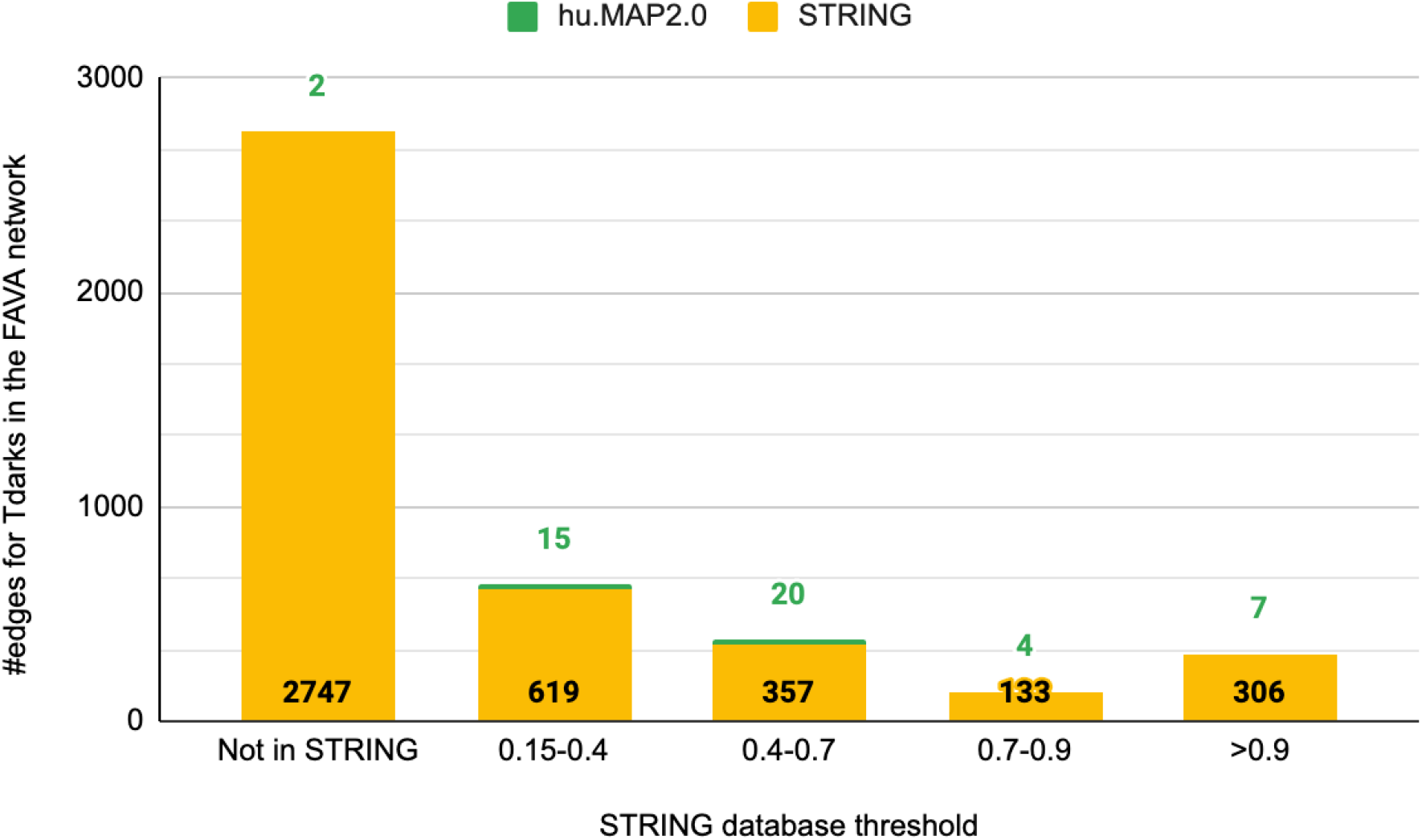
The Network A *.pdf* containing the network from Figure 5 with readable labels is provided.

**S6.** Characterization of the knowledge gain regarding understudied proteins. To assess what is the added value in terms of newly detected interactions by FAVA, we have compared the number of FAVA edges found in the STRING v11.5 databases and in hu.MAP 2.0. Four different score cutoffs were used for STRING (0.15 - includes all STRING edges, 0.4 - STRING medium confidence, 0.7 - STRING high confidence, 0.9 - STRING very high confidence). The STRING network was used for this comparison because it is one of the most comprehensive networks according to a comparison of 21 large-scale molecular interaction networks (10.1016/j.cels.2018.03.001). The majority of the FAVA interactions for Tdark proteins cannot be found in STRING at any score cutoff, and only very few interactions are in hu.MAP 2.0. This showcases both the added value from using co-expression data, but also the power of FAVA to detect these and further denotes the complementary nature of this network to current available experimental and literature knowledge.

**S7.** The Cytoscape session of the network and the analysis.

**Supplementary Table 1:** The table presents the results of a per-cluster enrichment analysis conducted on the FAVA network. This Table contains the full results of the enrichment analysis for each cluster (**Sheet 1 - Full functional enrichment results**). Additionally, a condensed version of the table is provided (**Sheet 2 - Selected terms of each cluster**), highlighting the most significant enriched terms in seven distinct categories: Pathways, UniProt Keywords, Cellular Compartments, Tissues, Biological Processes, Molecular Functions, and Diseases. These tables offer valuable insights into the functional characteristics of each cluster in the FAVA network.

## Notes

https://doi.org/10.5281/zenodo.6798182

https://www.proteinatlas.org/humanproteome/single+cell+type

https://www.ebi.ac.uk/pride/

https://doi.org/10.5281/zenodo.6803472

https://github.com/mikelkou/fava

### Competing Interest Statement

The authors have declared no competing interest.

### Summary of Updates

The revised manuscript showcases FAVA's performance that outperforms the state-of-the-art methods in coexpression analysis. FAVA has been rigorously tested on an extensive range of real-world data and meticulously benchmarked against numerous gold standards, attesting to its remarkable capabilities.

