## Supplementary figures and images for "FAVA: High-quality functional association networks inferred from scRNA-seq and proteomics data"

### Protein networks are commonly used for understanding how proteins interact. However, they are typically biased by data availability, favoring well-stu

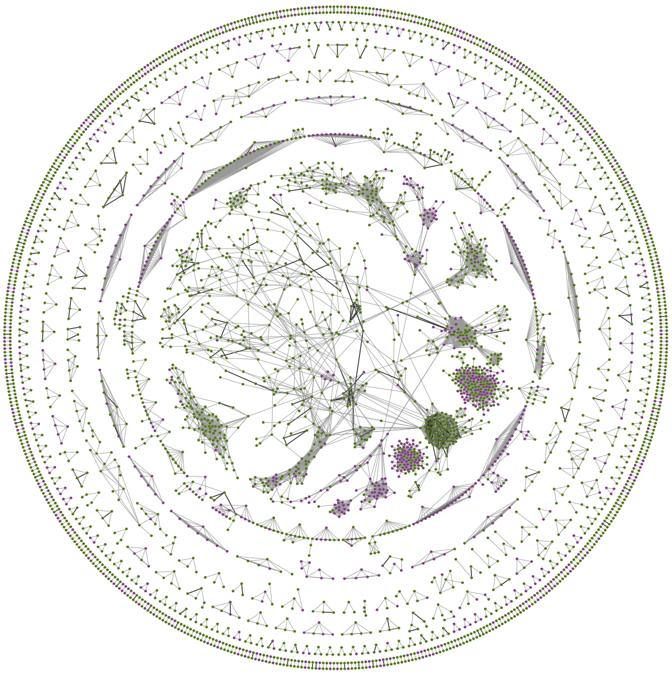
